# Behavioral and Modeling Evidence that Eye Movements Bias Self-motion Perception

**DOI:** 10.1101/2025.10.17.683136

**Authors:** Matt D. Anderson, Emily A. Cooper, Jorge Otero-Millan

## Abstract

To navigate the world, humans must integrate what they see with how they move. As the body moves, for example, the eyes rotate to explore the environment; these eye rotations, in turn, alter the visual signals used to judge body motion. Yet the practical impact of gaze dynamics on self-motion perception remains poorly understood. We tested how gaze position and gaze velocity shape self-motion perception—specifically heading direction—from visual signals in two behavioral experiments. In Experiment 1, we directly manipulated gaze position and velocity; in Experiment 2, a range of task demands evoked distinct gaze patterns. Across both experiments, heading estimates showed systematic, gaze-dependent errors: these estimates shifted toward the direction of eye motion and grew with horizontal gaze eccentricity and speed. A Bayesian ideal observer reproduced these error patterns across participants and tasks when it included three known features of visuomotor processing: (i) encoding retinal motion with eccentricity-dependent noise, (ii) underestimating eye-rotation speed, and (iii) a prior for moving straight ahead. These results reveal a lawful coupling between oculomotor behavior and heading perception. They further suggest that natural gaze strategies, such as keeping gaze near the optic-flow singularity and limiting pursuit speed, help mitigate gaze-dependent biases during navigation.

## Introduction

When we move through the world—whether walking, running, cycling, or driving—we use visual information to keep track of where we are going. One key piece of visual information is optic flow: the global pattern of motion signals cast across the visual field by the objects and surfaces around us. For an observer who travels in a straight line, there is a surprisingly simple way to determine the direction of self motion (heading) from optic flow: the optic flow pattern contains a focus of expansion, or ‘singularity’, and the location of this singularity corresponds precisely to the heading direction (Figure 1A). But there is a catch: this strategy only works if the observer does not rotate their eyes while they move. In controlled experiments that minimize eye movements, humans can indeed use the optic flow singularity to estimate heading (Gibson, 1950; Niehorster, 2021).

**Figure 1.**
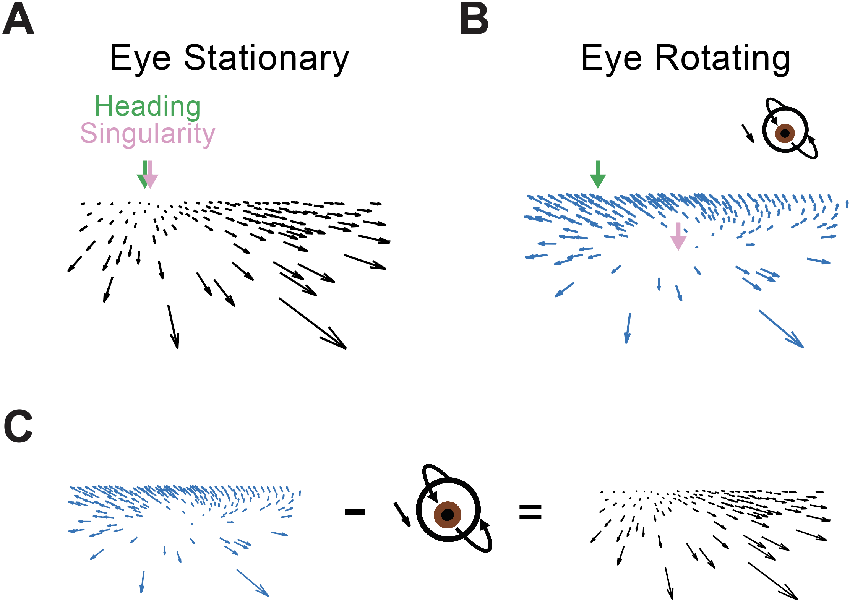
The effect of eye movements on optic flow signals. (A) When the eye is stationary in the head, and the observer translates in a straight line across a flat ground plane, the optic flow singularity (pink) corresponds exactly to the observer’s heading direction (green). Arrows are displaced for visibility. (B) Eye rotation adds a constant vector to all motion signals that comprise the optic flow field. This eliminates the one-to-one spatial mapping between the singularity and heading direction. (C) One simple computation that recovers the heading direction in the presence of eye rotation involves subtracting an estimate of the eye rotation from the retinal motion signal, and locating the singularity, as in panel A.

However, in the real world, humans continuously rotate their eyes, executing coordinated eye movements to track and fixate features like landmarks, obstacles, and signs while we navigate. These eye movements change the incoming optic flow signal, and remove the one-to-one mapping between the singularity location and heading direction (Figure 1B). However, since eye rotation adds a uniform motion vector to all signals in the optic flow, it is computationally possible to recover this mapping by subtracting the velocity of eye rotation from each of the retinal motion vectors that make up the optic flow field (Figure 1C). Research demonstrates that humans can recover heading during eye movements by estimating the velocity of their eye rotation with the help of extra-retinal cues such as efference copy signals or proprioceptive signals from the extraocular muscles (Banks et al., 1996; Bradley et al., 1996; Royden et al., 1992, 1994; Warren and Hannon, 1988). There are, however, costs to this process. Internal estimates of eye rotation are perturbed by sensory noise (Keller and Heinen, 1991), and systematic biases (Freeman and Banks, 1998; Freeman et al., 2010). These error sources suggest that gaze dynamics may impact the accuracy and precision of heading estimation. However, the contributions of eye movements to heading estimation errors, and the possibility that some eye movements are strategically deployed to mitigate these errors, are poorly understood (Angelaki and Hess, 2005; Matthis et al., 2022; Muller et al., 2023).

In this paper, we investigate these questions by manipulating, characterizing, and modeling how eye movements affect estimates of heading from optic flow. We focus on two features of eye movements that may reflect strategic adaptations to support heading estimation. First, when humans view optic flow they make intermittent saccadic eye movements that tend to position the center of gaze near to the singularity that reflects the heading direction (Chow et al., 2021; Knöll et al., 2018; Mei Chow and Spering, 2023; Niemann et al., 1999). We hypothesize that this singularity attraction reflects an adaptive strategy to minimize errors in heading perception, since heading discrimination is best when the singularity is placed in central (foveal) vision (Crowell and Banks, 1993; Warren and Kurtz, 1992). Second, when participants passively view optic flow, their eye movements tend to be slower than the foveal stimulus motion (Chow et al., 2021; Lappe et al., 1998; Niemann et al., 1999). Researchers have suggested that this quirk may reflect a strategy that balances the competing demands of heading estimation and other visual tasks requiring tracking of objects (Angelaki and Hess, 2005). For instance, slower eye movements minimize the amount of rotational compensation needed to recover heading, and in radially expanding optic flow fields produced by forward translation they also reduce the rate at which the fovea moves away the singularity. We therefore also hypothesize that slower eye movements support more accurate heading estimation. Our results support both hypotheses. More generally, our findings suggest that natural eye movement patterns during self motion are advantageous because they minimize gaze-dependent biases in heading estimation.

## Results

### Quantifying heading estimation performance as a function of gaze dynamics

In the first experiment, human participants (N=28) viewed optic flow stimuli that simulated forward locomotion at jogging speed across a flat ground plane in a random heading direction (from -15 to 15 degrees horizontally relative to head-center). While the optic flow stimuli were presented (Figure 2A), their gaze distance from the heading direction, and their gaze velocity were manipulated via a fixation target that varied in location and velocity on each trial (see Figure 2B-2C for all conditions). Gaze location was set to one of 12 positions relative to the true heading. Gaze velocity was manipulated by applying a multiplicative gain factor to the speed of the optic flow motion at the fixation target position. After stimulus presentation (1.25 seconds), participants used a mouse to slide a line horizontally to match their perceived heading direction. For full details, including catch trials in which gaze direction was also manipulated, see Methods section. On each trial, we computed the signed error as the angular deviation between the participant’s response and the true heading indicated by the optic flow. Positive errors indicated the response was more to the right than the actual heading.

**Figure 2.**
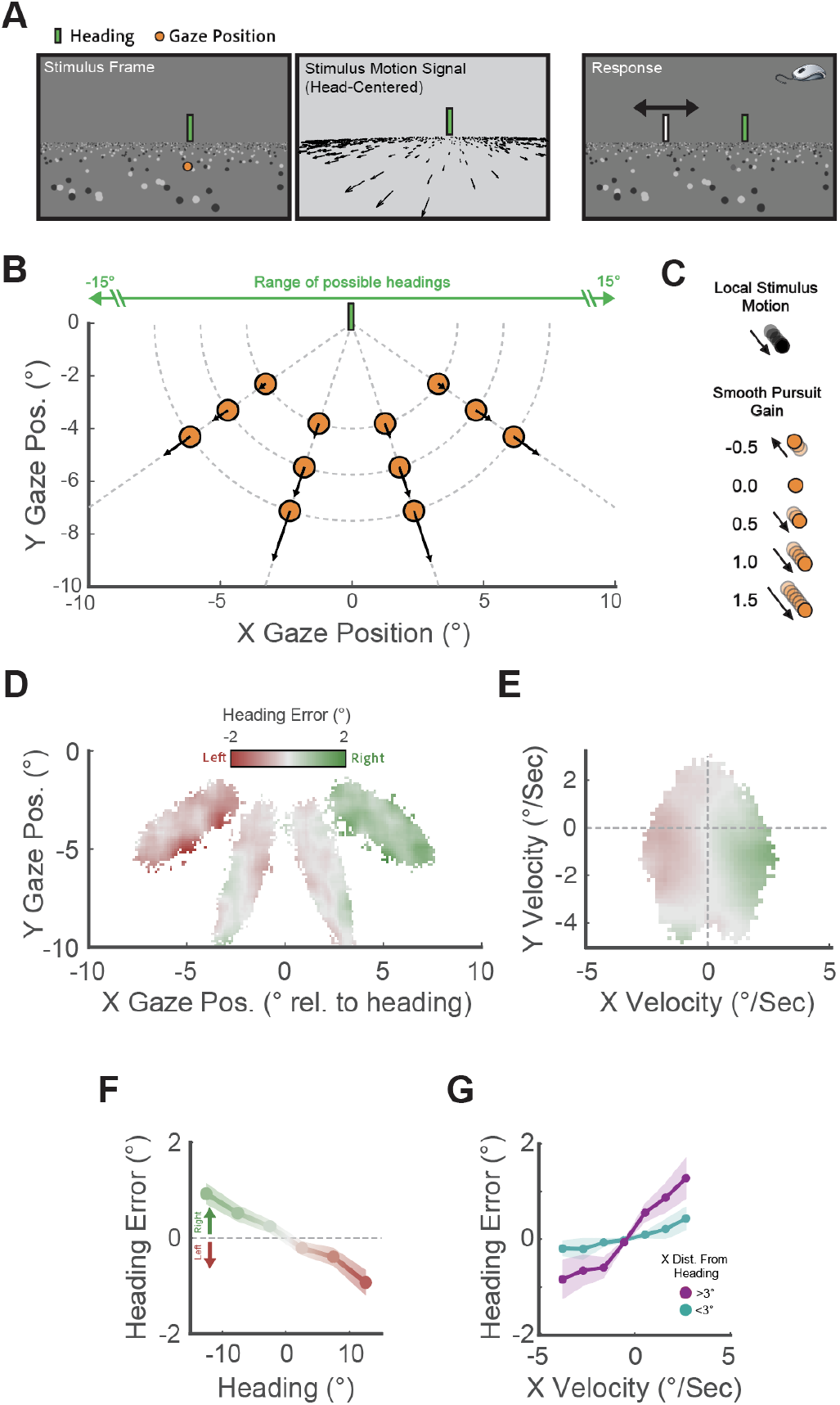
Experiment 1 methodology and results. (A) Example frame, motion signal, and response screen. After viewing the optic flow, participants responded by sliding the white line horizontally to match the perceived heading direction. The green line, indicating the true heading, was only shown to participants after they responded. The response screen showed a freeze-frame of the optic flow stimulus. (B) Orange dots show the 12 unique x-y gaze positions, specified by 4 meridians, and 3 radii. Gaze positions were defined relative to the true heading direction, which in this example is straight ahead (0 degrees). Arrows give the velocities of the optic flow stimulus at each location in degree/second. (C) The speed of the fixation point on each trial was modulated by a multiplicative gain factor applied to the instantaneous velocity of the optic flow at that location. Five different gains were tested. (D) Heading response errors as a function of x and y gaze position. Each pixel represents the sample-average. Pixels containing less than 5 participants’ data were removed, and data were smoothed with a gaussian kernel, *σ*=1 degree. (E) Heading response errors as a function of x and y gaze velocity. Data was smoothed in the same way as the position data. (F) Response errors as a function of true heading direction. Responses were binned based on 7 equally spaced headings spanning +-15 degrees. (G) The effect of horizontal gaze velocity (collapsing over vertical velocities) on heading errors varies with horizontal gaze distance from heading. Bins were equally sized with widths of 1 degree. Shaded error bars represent one standard error over participants.

### Heading errors vary with gaze distance from heading, gaze velocity, and heading direction

Figure 2D and E show the average heading errors as a function of gaze position (D) and velocity (E). Qualitatively, we observed that positive heading errors (corresponding to a rightward bias) were associated with both rightward gaze positions and rightward gaze velocities. Negative heading errors, corresponding to a leftward bias, were associated with both leftward gaze positions and leftward gaze velocities. Moreover, heading was biased overall towards the center of the screen/head, independent of gaze (Figure 2F). We fitted a linear mixed model to these errors with fixed effects for gaze position and velocity (separately for the horizontal and vertical dimensions), as well as heading direction. We limited interaction terms to two way interactions between gaze position and velocity in each x/y dimension (Table 1). Because heading direction in our experiment only varied horizontally, we hypothesized that gaze dynamics in the x dimension would be most associated with heading errors. A significant main effect of horizontal gaze velocity was modulated by an interaction with horizontal gaze position. In particular, higher horizontal gaze velocity was associated with a stronger heading bias in the direction of eye motion, and this bias increased with gaze positions farther to the left or right of the true heading direction (Figure 2G). We also found that the effect of heading direction, characterized by a bias towards the center of the head (Figure 2F), was statistically significant.

**Table 1.** Linear mixed model results from Experiment 1. Fixed effect coefficient estimates, standard errors (SE), degrees of freedom (DF), *t*-values, and *p*-values. Significant effects (*p <* 0.05) are indicated with bolding.

|  | Estimate | SE | DF | *t* | *p* |
| --- | --- | --- | --- | --- | --- |
| Gaze X Position (°) | 0.00 | 0.07 | 27.05 | 0.01 | 0.991 |
| Gaze Y Position (°) | 0.00 | 0.03 | 31.96 | 0.09 | 0.929 |
| **Gaze X Velocity (°/Sec)** | **0.37** | **0.14** | **26.67** | **2.56** | **0.016** |
| Gaze Y Velocity (°/Sec) | -0.03 | 0.07 | 7105.00 | -0.37 | 0.713 |
| **Heading Direction (°)** | **-0.07** | **0.00** | **9005.00** | **-22.45** | **<.001** |
| **Gaze X Position : Gaze X Velocity** | **0.06** | **0.03** | **8953.00** | **1.99** | **0.047** |
| Gaze Y Position : Gaze Y Velocity | 0.01 | 0.03 | 3076.00 | 0.33 | 0.744 |

Taken together, our data are consistent with the hypothesis that keeping eye movement speed slow and keeping gaze close to heading reduce errors in heading estimation. Moreover, the errors we observed were not random, but rather had a specific pattern: participants’ heading estimates were biased towards the direction of eye motion. However, these results do not speak to whether or not people deploy these eye movement patterns strategically during heading estimation. We address this question in a second experiment.

### Different tasks evoke different gaze strategies, which in turn affect heading errors

In a second experiment, participants (N=10, 5 of whom also participated in the previous experiment), viewed optic flow stimuli that simulated locomotion across a flat ground plane, this time consisting of green and red circles and squares. The heading direction always started straightahead, and then transitioned to a random heading direction (from -15 to 15 degrees horizontally relative to head-center) after a random delay of 1-3.5 seconds. There were three possible tasks: a heading estimation task, a visual search task, and a dual-task. The heading estimation task was the same as described previously. The visual search task required participants to determine the presence/absence of a specific target type, defined by a unique color-shape combination (Figure 3A). Participants were informed of the target type before each trial, and responded after stimulus presentation (like in the heading task). The dual task required participants to perform both the heading and visual search tasks on the same stimulus/trial. The order in which they made the heading or visual search responses was randomized. Trials were blocked by task, and participants were always informed of which task to perform before viewing the stimulus. Participants were free to move their eyes however they liked.

**Figure 3.**
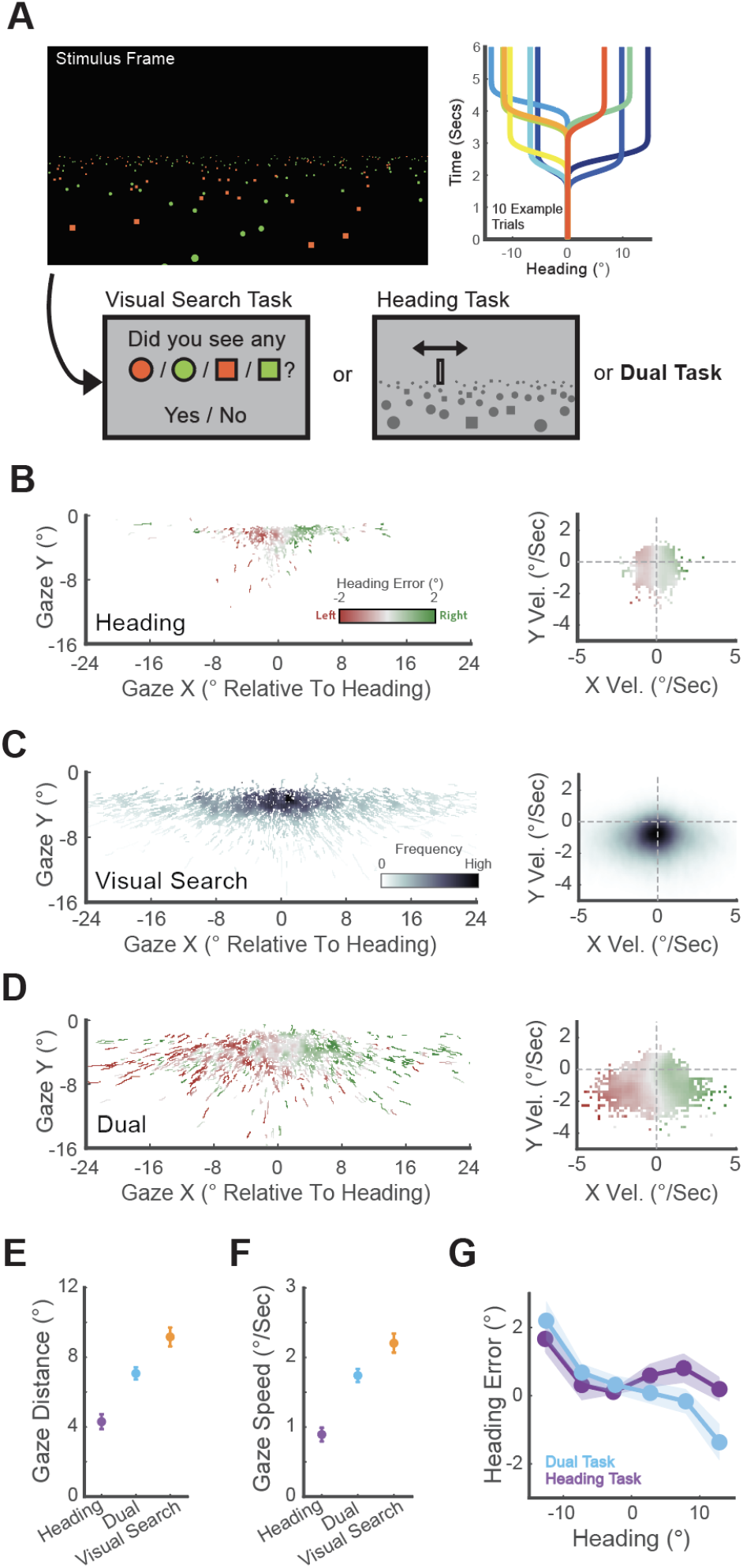
Experiment 2 methodology and behavioral results. (A) Example frame of the optic flow stimulus composed of red and green circles and squares. The graph (right) shows the temporal heading profiles of 10 example trials. Based on the presented stimulus, participants performed the search task, heading estimation task, or both (i.e., dual task). (B) Effect of gaze position and velocity on heading errors in the heading task. Data were smoothed with a gaussian kernel where *σ*=1 degree. (C) Heatmap of the gaze and velocity distribution in the visual search task (there are no heading errors since no heading responses were collected). (D) Effect of gaze position and velocity on heading errors in the dual task. Data were smoothed as in panel B. (E) Mean gaze distance from the heading direction split by task. (F) Mean gaze speed split by task. (G) Response errors as a function of heading, split by task. Error bars represent 1 standard error.

The task manipulation was effective at evoking different gaze strategies. In the heading task, participants tended to fixate stimulus locations near the heading direction (Figure 3B, left). Here, the shapes moved slowly, as did participants’ eye movements (Figure 3B, right). In the visual search task, participants fixated locations farther from the heading direction, where the shapes were larger and less crowded, but also moved faster (Figure 3C). In the dual task, they appeared to use a combination of these two strategies (Figure 3D). These trends were confirmed statistically: Distance of gaze from heading varied significantly between tasks (F(2,18) = 33.16, p <.001), and all follow-up pairwise comparisons were significant (p <.01), suggesting that participants fixated closest to heading in the heading task (mean=4.30 degrees, SE=0.42), followed by the dual task (mean=7.07 degrees, SE=0.34), and then the visual search task (mean=9.16 degrees, SE=0.54; Figure 3E). Gaze speed also varied between tasks (F(2,18) = 49.42, p = <.001), and all follow-up pairwise comparisons were significant (p <.01), suggesting that participants’ eye movements were slowest in the heading task (mean=.89 degrees/second, SE=0.10), followed by the dual task (mean=1.74 degrees/second, SE=0.09), and then the visual search task (mean=2.21 degrees/second, SE=0.14; Figure 3F). Thus, task demands affected where participants looked, and how fast the eyes moved. The heading estimation task was most associated with gaze positions near the heading and slow gaze speeds, consistent with our hypotheses that these gaze dynamics support heading estimation.

As before, we fitted a linear mixed model to the heading errors, collapsing over the heading and dual tasks (Table 2). One caveat of this analysis is that, because eye movements tended to track the expanding optic flow stimulus, horizontal gaze position and horizontal gaze velocity were more highly correlated than in the first experiment (r = 0.64 in this experiment versus r = 0.40 in the previous one), creating multicollinearity. This analysis revealed a significant effect of horizontal gaze distance from heading, and a significant effect of heading direction, characterized by a centerbias as in the first experiment (Figure 3G). Qualitatively, a horizontal velocity trend was observed (Figure 3A, 3C), but this was not borne out statistically, possibly due to predictor multicollinearity. Overall, these results replicate the key trends in the first experiment and suggest that humans may deploy gaze dynamics strategically to reduce heading estimation errors when heading is taskrelevant.

**Table 2.**
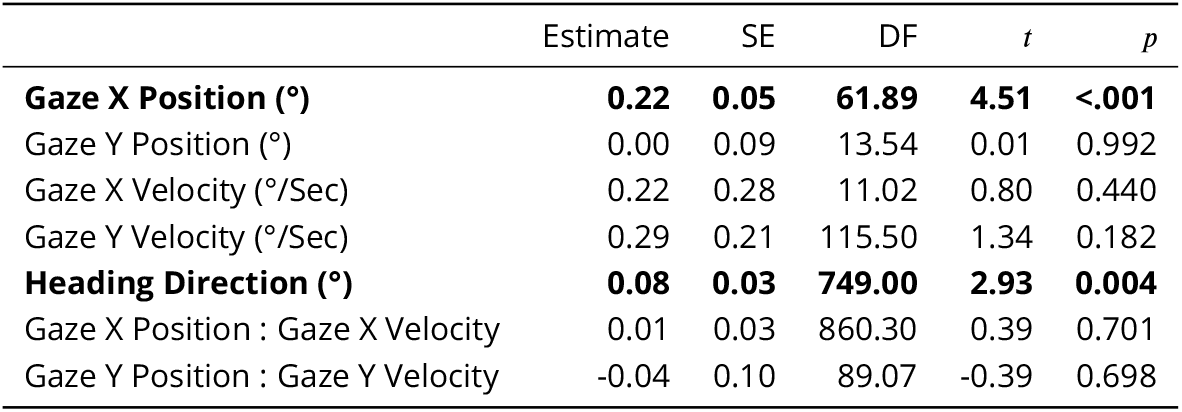
Linear mixed model results from Experiment 2. Fixed effect coefficient estimates, standard errors (SE), degrees of freedom (DF), *t*-values, and *p*-values. Significant effects (*p <* 0.05) are indicated with bolding.

| | Estimate | SE | DF | $t$ | $p$ |
| --- | --- | --- | --- | --- | --- |
| **Gaze X Position (°)** | **0.22** | **0.05** | **61.89** | **4.51** | **<.001** |
| Gaze Y Position (°) | 0.00 | 0.09 | 13.54 | 0.01 | 0.992 |
| Gaze X Velocity (°/Sec) | 0.22 | 0.28 | 11.02 | 0.80 | 0.440 |
| Gaze Y Velocity (°/Sec) | 0.29 | 0.21 | 115.50 | 1.34 | 0.182 |
| **Heading Direction (°)** | **0.08** | **0.03** | **749.00** | **2.93** | **0.004** |
| Gaze X Position : Gaze X Velocity | 0.01 | 0.03 | 860.30 | 0.39 | 0.701 |
| Gaze Y Position : Gaze Y Velocity | -0.04 | 0.10 | 89.07 | -0.39 | 0.698 |

### These heading errors are well-explained by an ideal observer model combining sensory noise and motion estimation biases

We found that the gaze-dependent heading estimation errors observed in our two experiments can be explained by three known properties of the visual system: eccentricity-dependent noise in retinal motion encoding (Crowell and Banks, 1996; McKee and Nakayama, 1984), biases in eye movement velocity estimation (Aubert, 1887; Freeman and Banks, 1998; Freeman et al., 2010), and biases in body/head orientation estimation (Cuturi and MacNeilage, 2013; Sun et al., 2020; Warren and Saunders, 1995; Xing and Saunders, 2016). Specifically, we formulated a Bayesian ideal observer model for heading estimation, fitted this model to trial-by-trial data from Experiment 1, and assessed the model’s ability to capture the observed pattern of heading errors in both experiments. The model observer encodes the optic flow stimuli presented to participants in the experiment, which is comprised of a set of dots with specific positions and velocities. These dots are projected onto a model observer’s retina based on eye tracker measurements of the participant’s gaze position and velocity (see Methods). The model observer then estimates their heading based on these retinal motion cues (Figure 4A).

**Figure 4.**
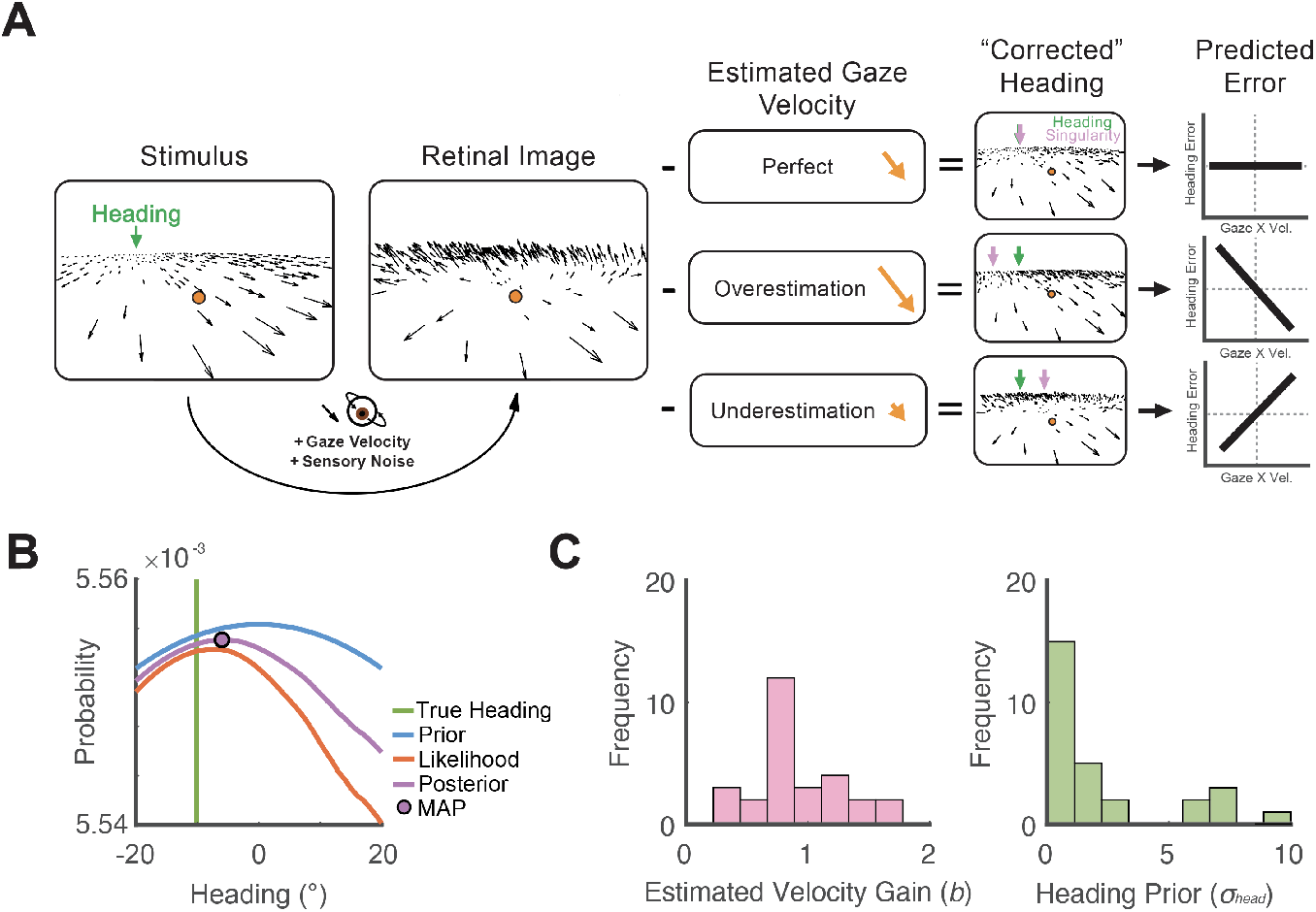
Bayesian ideal observer model fitted to participant data from Experiment 1. (A) The model’s retinal image is constructed by adding an eye velocity vector and eccentricity-dependent sensory noise to the stimulus input (that human observers saw). The model then compensates for eye rotation by subtracting the *estimated* eye velocity from the retinal input. Over and under estimation biases the singularity away and towards the direction of eye rotation respectively. The shift in the singularity produces a corresponding bias in heading perception. (B) An example trial illustrating how the model generates a heading response by computing the product of the center-prior and likelihood, over a range of heading directions, and selecting the heading that maximizes the posterior (MAP). (C) Distributions of the two fitted parameters: *b* and *σ*_*head*_, over the sample of 28 participants from Experiment 1.

In humans, retinal motion signals are perturbed by eccentricity-dependent sensory noise: noise is lowest near the fovea and highest in the periphery. This means that the precision with which an observer encodes stimulus velocity will depend on gaze position. We modelled this foveation using measurements of human motion sensitivity and spatial pooling data measured across the visual field (Figure 4A, left) (Crowell and Banks, 1996; McKee and Nakayama, 1984). Since detection of the singularity likely involves identifying a region of the retinal image with slow motion, it stands to reason that this eccentricity-dependent noise may impact heading estimates as seen in our data.

Using the noisy motion signals, the model then computes the posterior distribution of heading directions as the product of the likelihood and prior. The likelihood function is the probability of the noisy sensory input, given the expected signal associated with a hypothesized heading direction (i.e., an optic flow template). Critically, the likelihood function encodes these optic flow heading templates in a head-centered coordinate system. This means that the model must compensate for the eye movement velocity before computing the likelihood. Since the degree of compensation is unknown, we fit this parameter to each participants’ data as a multiplicative scaling constant (*b*) that quantifies the proportion of the actual eye velocity that is subtracted from the retinal signal before computing the likelihood. Previous work suggests that humans consistently underestimate the velocity of their smooth pursuit eye movements (*b <* 0) (Aubert, 1887; Freeman and Banks, 1998; Freeman et al., 2010), but there is little research on how this may impact heading perception. Qualitatively, an underestimation of eye velocity could explain the velocity-dependent heading error patterns we observed in our data as follows: if an observer uses eye velocity cues (such as extra-retinal signals) to compensate for eye velocity (see Figure 1), but underestimates the rate of eye rotation, the resulting optic flow signal contains a singularity, and inferred heading direction, that is biased *towards* the direction of eye motion (Figure 4A right).

Finally, the ideal observer also has a prior that reflects an assumption about which heading directions are most likely, in the absence of any sensory information. Based on previous work that demonstrates a strong heading bias towards the center of the head (Cuturi and MacNeilage, 2013; Sun et al., 2020; Warren and Saunders, 1995; Xing and Saunders, 2016), and our own data described above, we modeled this as a Gaussian with a mean of zero, and a standard deviation (*σ*_*head*_). This prior standard deviation was also fitted to each participant.

Figure 4C (left) shows the distribution of fitted gain parameters across our participant sample from Experiment 1. Although the distribution average is lower than 1 (mean =.95, median =.88), reflecting undercompensation for eye velocity, this effect was small and not statistically significant (*t*(27)=-.84, *p*=.408). The distribution of prior standard deviation parameters is highly positively skewed (Figure 4C, right), indicating that, consistent with previous work (Sun et al., 2020; Warren and Saunders, 1995; Xing and Saunders, 2016), heading perception is biased towards straight ahead.

The complete ideal observer model explained 16% of the variance in heading errors (Table 3). Expressed as a proportion of the maximum explainable variance, which adjusts for participant noise (see Methods), this model explained 71% of response variance (Table 3). Moreover, this model accurately reproduced the horizontal gaze position effect (Figure 5A), the horizontal velocity effect (Figure 5B), and the center-bias (Figure 5C). These findings suggest that our model predicts the measured patterns of human data well, and supports the notion that heading perception is biased by a combination of gaze dynamics and perceptual inferences.

**Table 3.** Bayesian ideal observer performance, fitted to data from Experiment 1. *R*^2^, noise-corrected *R*^2^ (see Methods), and mean squared prediction error (MSE) for different variants of the model, trained and tested via 10-fold cross-validation. In all model variants, the complete model was modified one parameter at a time.

| | $R^2$ | Noise-Corrected $R^2$ | MSE |
| --- | --- | --- | --- |
| Complete model | 0.16 | 0.71 | 7.77 |
| Uniform noise | 0.15 | 0.68 | 7.86 |
| Gaze velocity gain of 1 | 0.07 | 0.33 | 8.70 |
| Uniform prior | 0.10 | 0.46 | 8.70 |

**Figure 5.**
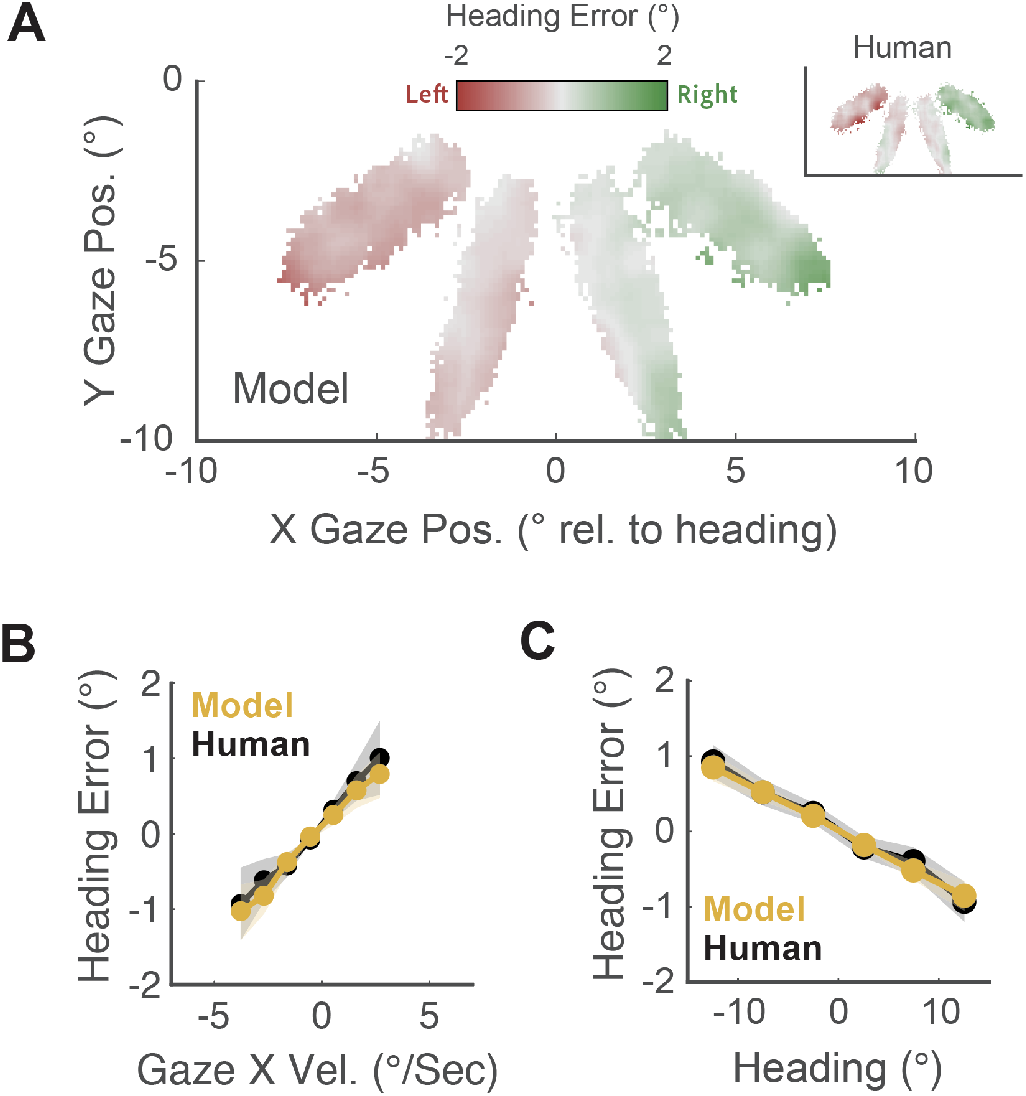
Ideal observer performance for predicting human heading estimation in Experiment 1. (A) The model generates heading errors as a function of horizontal gaze position in the same direction and magnitude as the human data (inset). Pixels containing fewer than 5 participants’ data were removed, and data were smoothed with a gaussian kernel where *σ*=1 degree. (B) Model and human heading errors are positively correlated with horizontal gaze velocity, reflecting a systematic bias towards the direction of eye movement velocity. Velocity bins were separated uniformly by 1 degree/second. (C) Model and human heading errors are negatively correlated with heading direction, reflecting a center-bias in heading estimates. Responses were binned into 7 heading directions linearly spanning +-15 degrees. Shaded error bars represent 1 standard error.

All three of the visual properties incorporated into our model (eccentricity-dependent retinal motion noise, biases in eye movement velocity estimation, and biases in body/head orientation estimation) contributed to the goodness of fit. Removing each property individually reduced the variance explained and increased the prediction error (Table 3).

### The ideal observer generalizes across tasks

The Bayesian ideal observer formulated for Experiment 1 can be applied to Experiment 2 with two small data processing steps. First, we isolated eye movements that occurred after the heading transition (1 to 3.5 seconds in duration). Second, we generated model responses for every smooth eye movement period within this epoch (excluding saccades), and averaged over all periods. This gives us a single model response for each trial, in a similar format to Experiment 1. For the 5 participants that participated in both experiments, we used the fitted parameters from Experiment 1, and for the remaining participants, we used the Experiment 1 averages. The model was not finetuned on any data from Experiment 2.

In Experiment 2, the model explained 18% of the variance in human errors, and 89% of the explainable variance in the heading task and the dual task. Splitting by task reveals that more variance is explained in the dual task (*R*^2^=.23) than the heading task (*R*^2^=.10). This may reflect an increase in both model and human response bias as a function of gaze speed, which is greater in the dual task than the heading task (Figures 6B). The model accurately reproduced the pattern of human heading biases associated with horizontal gaze position (Figure 6A), velocity (Figure 6B), and the bias towards straight-ahead (Figure 6C, although in this experiment, this bias was only notable for the dual task). Clearly, the systematic biases in heading perception we observed in Experiment 1 are replicable across different tasks and participants, and are well-captured by this model.

**Figure 6.**
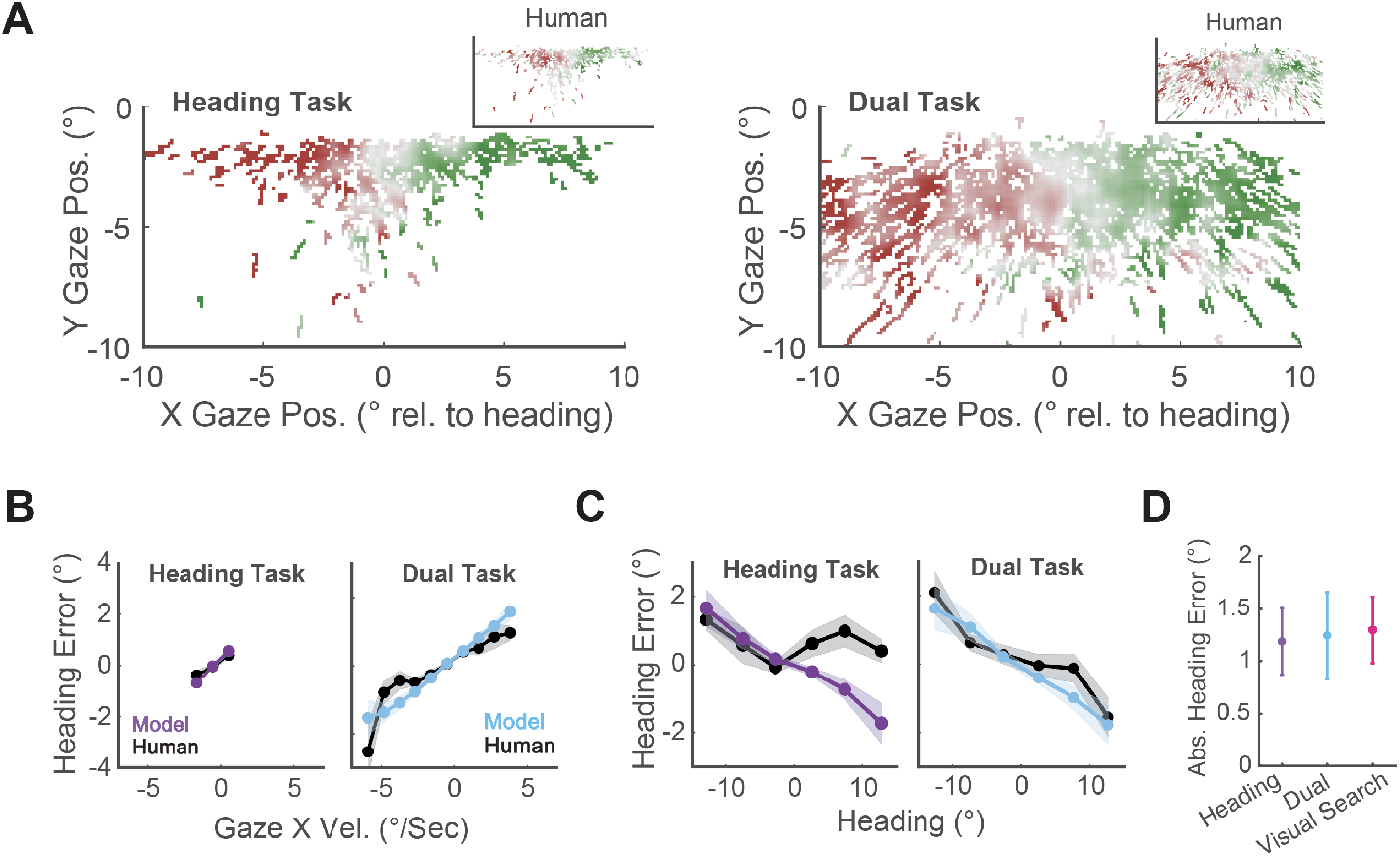
Model predictions for heading estimation errors in Experiment 2. (A) In both tasks, the model generates heading errors as a function of horizontal gaze position in the same direction and magnitude as the human data (insets). Data were smoothed with a gaussian kernel where *σ*=1 degree. (B) Model and human heading errors are positively correlated with horizontal gaze velocity, reflecting a systematic bias towards the direction of eye movement velocity. Data were binned into velocity bins 1 degree/second wide, and contained a minimum of 5 participants’ data. (C) Model and human heading errors are negatively correlated with heading direction, reflecting a center-bias in heading responses. Data were binned into 7 equally spaced bins spanning +/-15 degrees. (D) Model absolute (unsigned) heading error (in degrees), split by task. Error bars represent 1 standard error.

Thus far, we have characterized the contribution of different gaze strategies to heading perception. In the visual search task, participants chose to fixate stimulus locations farther from the heading direction. Based on this observation, if we could access the participants’ heading percept on these trials, what would it look like? Of course, we do not have heading responses from the visual search task, but our model gives us a unique opportunity to probe this question. We generated model responses in the same way as above, but used the human eye movement data from the visual search task. The resulting model-generated absolute response errors are shown in Figure 6D, alongside the other two tasks. Qualitatively, the model heading errors were in fact greatest in the search task, which suggests that tasks that require participants to fixate more eccentric, fastermoving targets can cause greater impairments in heading perception. However, this difference was small and not statistically significant.

## Discussion

### Effects of gaze position and velocity on heading estimation

We investigated how human gaze dynamics affect perceived heading during simulated self motion. Our results suggest that there is a reliable bias in perceived heading towards the direction of gaze position and motion. An important caveat is that in both of our experiments, gaze position (specifically gaze distance from heading) and gaze velocity were somewhat correlated – this is expected for eye movements that track elements an expanding optic flow field consistent with forward translation. In Experiment 1, singularity gaze distance appeared to modulate the effects of gaze velocity on heading estimation, but in Experiment 2 we did not find evidence for this modulation. Previous work has shown a strong tendency for people to put the center of gaze near the singularity in optic flow (Chow et al., 2021; Knöll et al., 2018; Mei Chow and Spering, 2023) and suggested that this behavior reflects an effort to place the most informative motion signals for heading direction discrimination in the central (foveal) retina (Crowell and Banks, 1993; Warren and Kurtz, 1992). Our results are thus consistent with these findings, and build on them by suggesting that the effects of singularity distance on heading estimation interact with gaze velocity.

We found that both the gaze position and velocity effects were most notable in the horizontal direction. This finding may also reflect the geometric properties of the stimulus - in our ground-plane stimulus, motion signals further from the singularity in the horizontal direction are more sensitive to noise perturbations for heading prediction than in the vertical direction (for discussion, see Crowell and Banks, 1993). However, this trend might also reflect a general property of heading perception, since most environments are characterized by a rigid ground plane (albeit with additional objects and vertical/oblique surfaces). The role of gaze position may also interact with motion anisotropies in visual discrimination. Humans have greatest sensitivity to horizontal and vertical motion directions (Ezzo et al., 2023; Gros et al., 1998). While we do not model these anisotropies in this paper, they have implications for predicting ‘optimal’ gaze behavior during self motion, and this represents a fruitful avenue for future research.

### The source of gaze-related heading errors

We propose that the visual mechanisms that underlie the effects of gaze dynamics on heading estimation include a retinal or extra-retinal signal that corrects for the effect of eye velocity on the optic flow signal, but underestimates the speed of eye rotation. Supporting this idea, a Bayesian ideal observer fitted to participants’ data, that incorporates this mechanism (among others) reproduces the pursuit bias described above, and predicts trial-to-trial heading errors well. This finding is consistent with a wealth of evidence that suggests humans tend to underestimate the speed of moving stimuli (Stocker and Simoncelli, 2006; Weiss et al., 2002; Welchman et al., 2008), and their own eye movements (Freeman and Banks, 1998; Freeman et al., 2010). Since most objects in the world, from a static observer’s perspective, move at slow speeds (Dong and Atick, 1995), underestimation might reflect an optimal tuning to natural motion statistics. How well this applies to the optic flow motion statistics created by self-motion is unclear, however. Recent work suggests that during natural self motion, gaze is rarely stable relative to the head, and objects only move at slow speeds on a small region of the foveal retina (Matthis et al., 2022; Muller et al., 2023).

Ultimately, the fitted gains for eye velocity estimation were only slightly less than 1 (and sta-tistically non-significant), and the associated human and model heading estimation biases were similarly small in terms of visual degrees (averaging 1.5 - 2 degrees for self-motion speeds of 3 degrees/second). However, the magnitude of underestimation also varies with the translational velocity of self motion. For example, our model predicts that, for an observer with parameters taken from the sample averages who is driving a car at 45 miles/hour, pursuing an object 15 degrees from the head will result in a heading error of approximately 10 degrees. This magnitude of error would be highly consequential in real-world driving scenarios.

Another important component of our model was the center-bias for heading estimation. Other studies have observed similar biases (Sun et al., 2020; Warren and Saunders, 1995; Xing and Saunders, 2016). In our model, this center-bias is embodied primarily as a straight ahead prior for heading. Replacing this with a uniform prior resulted in substantially worse predictions of human heading responses. However, because gaze position and velocity also bias heading, the center bias may also reflect a tendency for gaze statistics towards straight ahead. Theoretically, this center-bias could reflect a perceptual or response-based pattern, since the orientation of the head always corresponded to the center of the screen (so a reasonable response with no visual information is the center of the screen, given the uniform distribution of headings). However, there is some evidence that this bias is perceptual. For example, other tasks produce a similar behavior: in a 2AFC task where participants judged whether they were heading to the left or right of a visual probe, Warren and Saunders (1995) researchers also observed a center bias. Building on this insight, Xing and Saunders (2016) report that this bias is not affected by the distribution of headings in the experiment, and follows changes in head orientation.

One additional variable relevant to predicting human heading behavior, albeit to a lesser extent than the two mechanisms above (see Table 3), is eccentricity-dependent noise. Specifically, human motion sensitivity decreases with increasing distance from the fovea (Crowell and Banks, 1996). Modeling this non-uniformity with human measurements of motion speed and direction noise across the retina results in slightly better model performance than assuming foveal noise across the whole visual field. Since eccentricity-dependent noise effectively varies the relative weight given to the center-prior depending on the gaze position relative to the singularity (where feature density and informativeness regarding heading is greatest), this finding suggests that, in a typical Bayesian fashion, humans increasingly rely on priors when sensory information is noisier.

### Multitasking during navigation

In controlled optic flow experiments, the participant is typically given only one task: judge some visual or latent property of the stimulus. In the natural world, self motion is accompanied by multiple demands that humans have to juggle simultaneously. For example, as we walk through a city street, we may look out for cars, trip hazards, look at our phone to check a message, and pick a face out of a crowd, all while *simultaneously* monitoring our heading direction. Several recent studies have explored natural gaze dynamics during real-world locomotion, and they have revealed an interesting pattern: humans spend much of their time fixating nearby objects like future footholds, and intermittently switch their gaze to near-horizon locations that roughly correspond to their heading direction (Matthis et al., 2022; Muller et al., 2023). Our study replicates this behavior in a controlled lab setting: In our dual task experiment, where participants simultaneously judged the presence/absence of targets and their heading direction, we observed a gaze distribution that suggests participants were both fixating targets towards the bottom of the visual field, and the singularity at the middle of the visual field (Figure 3). This finding contributes to growing evidence that humans actively sample visual information from different regions of the visual field, to jointly optimize performance for multiple tasks during self motion.

When viewing optic flow stimuli, humans also perform eye movements to stabilize moving features on the retina. One quirk of the stabilizing eye movement is that it often undercompensates for object motion, resulting in a tracking gain of <1. We tested a leading explanation of this phenomenon: that slow pursuit optimizes heading perception during concurrent visual tasks (Angelaki and Hess, 2005). Importantly, the magnitude of the heading estimation errors observed in our study scaled approximately linearly with the horizontal speed of pursuit, suggesting that slow pursuit is indeed advantageous to heading perception. In situations where gaze is drawn away from the singularity, such as during our visual search and dual task experiments, the visual system has little choice but to (at least partially) track visual objects. While our findings provide little evidence that, in these situations, the visual system actively modulates pursuit gain, we do find that the eye movements that accompany the gaze distributions in these tasks are associated with stronger biases in perceived heading. That is to say there is a cost associated with multitasking, and slow pursuit may be a method of minimizing this cost.

### Caveats and limitations

An important limitation is that in all our experiments, head orientation was fixed using a chin and head rest. In natural tasks with no biomechanical constraints, humans typically coordinate head, eye, and trunk movements to minimize eccentric gaze positions, and keep the eyes facing forward (Burlingham et al., 2024; Foulsham et al., 2011; Land, 2004). Gaze eccentricities in both experiments were not so extreme as to be atypical of normal viewing conditions (i.e., within 15 degrees of head-center, Burlingham et al., 2024; Foulsham et al., 2011), but the contribution of head/trunk rotations, their modulation of gaze effects, and the additional computations required to discount these rotations from the optic flow signal, to recover heading direction, remain unclear. We also cannot determine the relative contributions of retinal and extra-retinal signals for the estimation of eye velocity. This experiment was not conducted in absolute darkness, so spatial information such as the motion of the screen’s edges, could be used to estimate eye velocity independently of efference copies or proprioception. Future studies could compare the errors produced in the same task with and without various visual and non-visual cues for eye rotation.

An additional limiting factor that we do not account for in this study is the contribution of different sensory systems that ordinarily support self motion perception, such as vestibular sensation and proprioception. The vestibular organs are sensitive to linear acceleration cues that accompany locomotion, and proprioception refers to the set of internal signals that encode the position and orientation of our musculoskeletal system as it moves through space. Information from both these systems are integrated with visual signals during self motion (for reviews, see Fetsch et al., 2010 and Israël and Warren, 2005). In our experiments, the measured heading biases reflect scenarios where the vestibular and proprioception modalities give conflicting information relative to the visual modality, and it remains possible that, in natural environments, these extra-retinal systems may help the observer to compensate for errors caused by gaze dynamics. An interesting research idea for future work would be to employ a closed-loop design where stimulus motion is manipulated as the participant is actively moving through an environment. Measuring steering behavior, postural sway, and other responses as a function of their gaze dynamics, will give novel insight into how the senses work together during self motion to monitor and correct for online heading errors.

## Conclusions

Humans continuously move their eyes as they navigate through the natural world. Using behavioral data from two experiments, and a computational model, we investigated the contribution of these gaze dynamics on errors in heading estimation. We found that the position and velocity of gaze varies with task, and produces predictable errors in heading that can be explained by three core mechanisms: eccentricity-dependent retinal noise, underestimation of eye velocity, and a center heading bias. These findings shed new light on the causes of heading errors, and constitute a step towards understanding ways to mitigate their negative effects in real-world scenarios.

## Methods

### Participants

Twenty-eight participants were recruited in Experiment 1 (Mean age = 24.8 years, stdev. = 5.3; 22 females, 6 males). Sample size was determined by conducting a power analysis in G*Power on a pilot sample of 5 participants. Ten participants were recruited in Experiment 2, 5 of whom were also in Experiment 1 (Mean age = 30.2 years, stdev. = 4.8; 6 females, 4 males). In both experiments, participants were screened for normal visual acuity (all 20/30 binocular or better) and color vision (all completed 13/13 plates on the Ishihara test). No participants were removed in either experiment based on abnormal screening measurements. Both experiments were approved by the University of California, Berkeley, Institutional Review Board. All participants provided informed consent and were compensated for their time.

### Stimuli and Tasks

In both experiments, binocular gaze position was recorded using an Eyelink 1000+ at a sampling rate of 500 Hz calibrated using a 9-point calibration grid. Participants viewed stimuli while seated at an eye-height of 1.23 meters, and their head was stabilized with a chin and forehead rest. The experiment was performed in a dimly lit room.

In a series of trials, stimuli were presented on a large computer monitor (Samsung S95C 77” TV), that subtended 66.36 by 40.53 degrees, at a viewing distance of 1.3 meters. Each stimulus consisted of 1000-2000 shapes scattered randomly in world-space across a flat ground plane, and projected onto the screen plane using perspective projection. The far clipping plane was 60 meters from the viewing position. We generated optic flow by simulating forward translation through space, parallel to the ground plane, at jogging speed (3.25 meters/sec; Experiment 2 used an additional running speed of 5.76 meters/sec, but we collapsed over speeds after finding no effect on behavior). On-screen shape locations were updated on each frame (at 144 Hz). In both experiments, heading directions were sampled from a uniform distribution with a range of -15 degrees (leftward) to 15 degrees (rightward). Shapes had a virtual diameter of 5.4 cm, and varied in projected size based on perspective projection. Shapes had a limited lifetime of 1.25 or 1 seconds (for Experiment 1 and 2, respectively), and expired shapes were repositioned to new random world locations. To avoid all shapes repositioning at once, shapes were initialized at a random time within their lifetime at the start of each trial.

#### Experiment 1

In this experiment, we used a moving fixation point to manipulate the velocity and location of gaze while viewing optic flow. Prior to the optic flow stimulus onset, a single yellow fixation dot appeared at a pre-determined location and moved with a pre-determined motion trajectory for 0.75 seconds. The optic flow stimulus then appeared for 1.25 seconds, during which time the fixation dot continued on its trajectory. The optic flow shapes consisted of black and white circular dots against a mid-gray background. On a given trial, the fixation dot assumed one of 12 unique locations relative to the simulated heading (see Figure 2B). Locations were specified by 4 meridians (215, 252, 288, 325 counter-clockwise angular degrees), and 3 radii along these meridians (4, 5.75, and 7.5 degrees of visual angle). For each fixation location, there were 5 unique gain conditions: -.5, 0,.5, 1, and 1.5. The gain refers to a multiplicative scaling factor that varied the speed of the fixation point relative to the underlying optic flow. Gain values of >1 indicate that gaze was moving faster than the optic flow, a zero gain represents stationary fixation, and negative gains reverse the gaze motion direction. The motion of the fixation point was computed by calculating the instantaneous (frame-to-frame) on-screen velocity, specified by its location on the ground plane, and the trial’s heading direction, and scaling this velocity by the gain. That is to say that the gain is applied to the projected velocity, rather than the world velocity. To ensure that our gain manipulation did not systematically modulate mean gaze position, we varied the initial location of the fixation dot such that, during stimulus presentation, the average gaze location was the same across all gain conditions.

To prevent participants from using motion afterimages to perform the task (reported during piloting), a motion mask was constructed by sampling dot sizes and speeds from the stimulus distribution, and rendering them at random positions and motion directions on the screen. Mask presentation was 0.3 seconds. After mask presentation, participants were shown a freeze-frame of the stimulus. A vertical line was rendered at the horizon of the scene, and participants adjusted its horizontal position using a mouse to match their perceived heading direction. Participants clicked a mouse button to submit their response, and feedback was given on each trial in the form of a green line that specified the true heading. Response time was unlimited, and no participants reported difficulty understanding the task.

Note that, since the fixation point motion followed the same geometry as the stimulus, participants could theoretically judge heading direction by simply locating the point where the path of the fixation point intersects with the horizon. To ensure that participants were in fact encoding the global properties of the stimulus, 25% of trials were ‘catch trials’, where the heading direction specified by the motion of the fixation point was randomly offset by +-15 degrees. An analysis of catch trials revealed that participant responses were substantially more correlated with the stimulus heading direction (*R*^2^=.81), than the fixation point motion direction (*R*^2^=.64). This confirms that participants were encoding global optic flow information in this experiment. These catch trials were removed for all reported analyses.

The experiment contained 480 trials per participant, split equally into 8 60-trial blocks. Conditions were randomly shu?ed across blocks. The eye tracker was recalibrated at the beginning of each block. The experiment lasted an average of 69.7 minutes (stdev. = 22.5 minutes).

#### Experiment 2

In this experiment, gaze dynamics were manipulated by giving participants different tasks to perform (Figure 3A). Each trial started with a fixation cross at the center of the screen. Participants clicked a mouse button to reveal the stimulus when they were ready. During stimulus presentation, the cross disappeared, and participants were free to move their eyes however they liked. The optic flow shapes consisted of red and green circles and squares against a black background. The entire stimulus was displayed for 6 seconds, and underwent three heading phases. In the initial phase, which lasted between 1 and 3.5 seconds (randomized between trials), the heading was fixed at straight-ahead. In the second phase, the heading transitioned from straight-ahead to a new random heading direction (within +/-15 degrees), over the span of 1.5 seconds. The temporal transition profile was a cumulative gaussian with a standard deviation of 0.21 seconds. In the third and final phase, which lasted between 1 to 3.5 seconds, the heading direction was fixed until the end of the trial. We included two simulated locomotion speeds: 3.25 and 5.76 meters/second, but collapsed across them in all reported analyses (after finding no effects/interactions). On each trial, a subset of shapes (18 exactly) were defined as ‘targets’, and were visible for a minimum of 40% of the trial duration (2.4 seconds). This was to ensure that the target locations in the visual search task (see below) were unpredictable, but always visible.

The critical manipulation in Experiment 2 was the task type. The experiment was split into six 30-trial blocks, and in each block participants performed a different task. There were three tasks: a heading estimation task, visual search task, and a dual task. The heading estimation task was exactly as in Experiment 1, except before stimulus onset the participant was given the verbal prompt “Estimate the heading”. The visual search task required the participant to determine the presence/absence of a pre-determined target. The target was defined by a unique color-shape combination (e.g., red circles or green squares). Since the target was defined by two feature dimensions (color *and* shape), it required the participant to effortfully search across the stimulus, and fixate different shapes in a serial manner (Treisman and Gelade, 1980; Wolfe, 2020). The participant was informed of the target type prior to each trial in the form of a verbal prompt that read, for example: “Search for the green squares”. After stimulus presentation, they made a present/absent judgment using two mouse buttons. Feedback was given on every trial in the form of a correct/incorrect tone. The dual task required the participant to judge both the heading and target presence/absence. Prior to stimulus onset, the prompt read, for example “Search for the red circles, and estimate the heading”. As in the two other tasks, participants only responded after the stimulus was presented. The order in which they responded was randomized between trials (and not known to the participant in advance). In all three tasks, participants were given no instructions regarding how they should move their eyes based on the task. The stimulus properties were the same across all three tasks. The experiment lasted an average of 56.0 minutes (std = 11.1 minutes).

The visual search task was designed to be sufficiently difficult to impact human gaze dynamics, but not so difficult as to render multitasking (joint performance of the heading task) impossible. An analysis of the proportion-correct data (true-positive and true-negative responses) revealed that the visual search (mean = 0.69, SE = 0.01) and dual task (mean = 0.67, SE = 0.01) both hit struck a balance between floor and ceiling performance. The dual task was slightly more difficult, but the difference was non-significant (*t*(9)=-0.81, *p*=0.44). We focus on heading performance in all analyses in the main text.

### Analysis

#### Data Cleaning

We took several steps to clean and parse the eye movement data to prepare it for visualization and statistical analysis. These processes were applied to the data from both experiments (where applicable). Since our main interest lies in the dynamics of slow, non-saccadic eye movements often called smooth pursuits, we first isolated the smooth phases. To do this, we first applied pupil area thresholds to remove blinks, and position, velocity, and acceleration thresholds to remove other eye movement artifacts. To segment smooth pursuits and saccades, we applied the Engbert and Kliegl (2003) saccade detection algorithm (*λ*=6). Saccadic eye movements were then removed from the data. Raw gaze position data were used without filtering where applicable, but to estimate gaze velocity, position data was first filtered with a Sovitzky-Golay filter, with a 1 degree polynomial, and 31-sample window.

Exclusion criteria were determined prior to statistical analysis and modelling. Entire trials in Experiment 1 were removed if: the smooth pursuit period during stimulus presentation did not exceed 0.8 seconds (over all periods), fixation position deviated from the target by an average of more than 3 degrees, the measured gain based on the eye movement data deviated from the gain of the fixation point by +/-2 (additional outliers were removed via a participant-specific threshold of mean +/-3 standard deviations), any saccade amplitude exceeded 4 degrees, saccade frequency exceeded 4 over the trial duration (3.2 per second), response time exceeded 10 seconds, and absolute heading error exceeded 15 degrees. Experiment 2 did not constrain participants’ eye movements, so only a subset of these exclusion criteria were applied (excluding saccadic criteria, and the target-distance criterion). In Experiment 1, these criteria led to the removal of 10.96% of trials, and in Experiment 2, 8.17%. In all analyzed eye movement data, we averaged gaze positions over the two eyes to improve the signal to noise ratio.

#### Statistical Testing

On each trial, we computed the heading estimation error as the signed angular difference between the true heading from the optic flow and the participant’s response. To test the statistical significance of our experimental manipulations, we fitted linear mixed models to these error data in R using the LME4 package (Bates et al., 2015). The random effects included participant intercepts, and participant-specific slopes for X position, X velocity, and Y position. Any additional random effects led to convergence issues, and all alternative permutations of models with maximal random effects generated the same set of significant/non-significant effects. Fixed effects were specified based on prior hypotheses, and included main effects of: X gaze position, Y gaze position, X gaze velocity, Y gaze velocity, and 2-way interactions between X position and velocity, and Y position and velocity. Parameters were fitted via maximum likelihood estimation, and statistical significance was estimated using Satterthwaite’s method of approximating the effective degrees of freedom (lmerTest package, Kuznetsova et al., 2017; note that this results in non-integer degrees of freedom in reported tables).

### Bayes Ideal Observer

#### Optic flow signal

We used a right handed world coordinate system in which the *x* axis is positive away, the *y* axis is positive leftward, and the *z* axis is positive up. We simulated the optic flow seen by an observer translating through this world parallel to a horizontal ground plane (i.e., in the positive *x* direction). The origin of the observer’s head coordinates was centered on the world origin but shifted upward by the observer’s simulated eye height: [0 0 *z*_*height*_]. The observer always started at this origin and then moved with a heading azimuth *θ*_*head*_ at a constant speed of *s*_*head*_. The self-motion velocity in head-centered coordinates **v**_**head**_ was therefore given by:

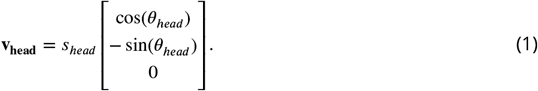

The associated optic flow was projected to a theoretical cyclopean eye located at the center of the head. As such, our ideal observer does not have access to stereoscopic depth information. The initial direction of the eye’s gaze was manipulated by specifying the azimuth (*θ*_*eye*_) and elevation (*α*_*eye*_) of the fixation point relative to straight ahead. This gaze direction was used to define the rotation matrix *R*_*eye*_ corresponding with a rotation of angle 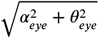 around an axis of rotation contained in the *yz* plane deviated by an angle tan^−1^(*α*_*eye*_, *θ*_*eye*_) from the *y* axis, yielding the eye’s translational velocity in head-coordinates as:

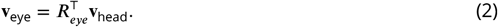

That is, the eye translates with the speed of the head, but the direction in eye coordinates is rotated according to the initial angle of gaze.

To determine the eye’s rotational (angular) velocity, we assert that the observer aims to stabilize the object at the fixation point *θ*_*eye*_, *α*_*eye*_ as it translates. We computed the angular velocity of the eye as the one produces no net flow at the point of fixation. The eye angular velocity is therefore proportional to the flow at fixation produced by translation (this motion vector is denoted by **f**_**t**_, with horizontal and vertical components *f*_*u*_ and *f*_*v*_) and can be computed according to the equation:

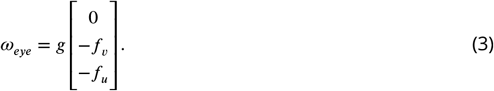

Intuitively this equation tells us that the eye needs to rotate around the y-axis to compensate for the horizontal flow and around the x-axis to compensate for vertical flow. Here, *g* denotes a multiplicative gain factor that determines whether the eye rotation is faster, slower, or the same as needed to maintain fixation. Negative gains indicate that the eye is moving in the opposite direction. Note that there are multiple angular velocities that would produce the same result with different torsional velocities. Here we adopted the solution with zero angular velocity around the gaze direction (*x*) axis.

Using the translation and rotation of the eye, we determined the ground truth optic flow for a set of *n* randomly distributed ‘world’ points on the ground plane using perspective projection. This optic flow signal comprises an *n* × 2 matrix of 2D flow velocities (**f**), denoted *F*. Each flow velocity in the matrix is associated with a particular angular retinal coordinate relative to the center of gaze (*θ*_*flow*_, *α*_*flow*_). The flow vectors, and therefore the flow matrix, can be decomposed into the sum of the flow due to translation and rotation (**f** = **f**_**t**_ + **f**_**r**_; *F* = *F*_*t*_ + *F*_*r*_); see Figure 1. Ultimately, *F*_*t*_ is the optic flow signal that indicates heading.

#### Ideal observer model for heading from optic flow

The motion samples that the ideal observer has access to are corrupted by measurement noise. The amount of measurement noise for visual motion is not always the same: it varies depending on the properties of the stimulus, such as the stimulus speed and retinal eccentricity (Crowell and Banks, 1996). We were particularly interested in modeling how retinal eccentricity-dependent changes in measurement noise might affect heading estimation. Thus, we relied on a previous psychophysical study that characterized measurement noise for single dot stimuli moving at different speeds and falling on different retinal eccentricities (Crowell and Banks, 1996). This study reports estimates of the standard deviations of the noise distributions for local motion in terms of speed and motion direction. We assumed that the measurement noise for each stimulus motion was unbiased. Specifically, we modeled the probability distributions of speed and direction measurements at each retinal eccentricity 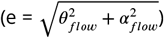 as a Gaussian centered on the ground truth speed and direction associated with the optic flow signal *F* and with standard deviations for speed (*σ*_*s*_(*e*)) and motion direction (*σ*_*ν*_(*e*)).

In addition to motion sensitivity varying by retinal eccentricity we also know that spatial sampling of visual information varies across the visual field. Points further from the center of gaze (or fovea) are increasingly downsampled based on front-end mechanisms such as photoreceptor and ganglion cell spacing on the retina (Rossi and Roorda, 2010), and downstream mechanisms, like cortical receptive fields (Anderson and Burr, 1987; Burr et al., 1998; Duffy and Wurtz, 1991). This downsampling likely contributes to the eccentricity-dependent changes in local motion sensitivity, thereby affecting optic flow encoding. In our main analysis, we thus used motion-sensitive receptive field sizes measured psychophysically by McKee and Nakayama (1984) and linearly averaged the vectors in each receptive field to generate a single pooled motion vector. We then applied the noise model described above to the pooled motion vector. This pooling technique was selected from a range of candidates based on the analysis described below in “Eccentricity-based downsampling.” We denote the resulting noisy measurement of flow as 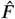.

Previous work suggests that the visual system can discount the effects of eye rotation on 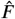 in estimates of heading from optic flow (Banks et al., 1996; Bradley et al., 1996; Royden et al., 1992, 1994; Warren and Hannon, 1988). That is, heading (*θ*_*head*_) is obtained by taking a measurement of *ω*_*eye*_ (derived from retinal and extra-retinal cues) and effectively subtracting the associated rotational flow matrix from the optic flow on the retina. Unlike local visual motion estimates, here we assume that the estimate of *ω*_*eye*_ is biased. Specifically, that the rotational speed may be underestimated (Freeman and Banks, 1998; Freeman et al., 2010). For simplicity, we do not model the variance of the underestimates. We thus assume that 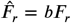 where *b* denotes the underestimation (or overestimation) of eye rotation speed. The resulting noisy measurement of optic flow due to translation 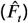 is modeled as follows:

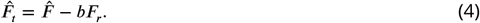

For simplicity, we also assume that the model observer’s estimate of pursuit *direction* is unbiased and noiseless. Our ideal observer then uses Bayes Rule to estimate their heading (*θ*_*head*_) from 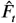. Specifically, they seek find the value of *θ*_*head*_ that maximizes the posterior probability 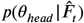 as determined by:

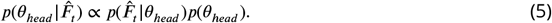

The likelihood function 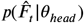 is determined by the noise properties of the encoder that we obtained from previous measurements (Crowell and Banks, 1996). This amounts to an assumption that the model has access to the noise distributions in the sensory inputs.

For a given 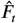, we solved for the likelihood numerically, by iterating over a range of possible headings (from -25 and 25 degrees in steps of 1 degree) and calculating the conditional probability under the assumption that each motion speed and direction estimate is independent.

The prior over heading (*p*(*θ*_*head*_)) reflects the observer’s prior assumptions about which heading directions are more likely than others. We modeled this as a Gaussian distribution with a standard deviation parameter *σ*_*head*_ and mean of 0 degrees, which is motivated by previous research indicating that human heading perception is biased towards the center of the head (Sun et al., 2020; Warren and Saunders, 1995; Xing and Saunders, 2016).

We assume that the observer’s perceived heading 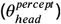 is the heading that maximizes the posterior over the range of tested headings. As such, our main ideal observer includes two free parameters and one fixed parameter that can introduce biases in heading estimations. The two free parameters are the gain factor on eye rotation speed *b* and a prior assumption that heading is straight ahead (embodied by the free parameter *σ*_*head*_). The eccentricity dependent noise parameter (*σ*_*s*_(*e*), *σ*_*n*_*u*(*e*)) was modeled based on prior data and was not included as a free parameter.

#### Fitting the model to the data

We assumed that the human participants’ heading responses reflect a direct read-out of their perceived heading on each trial. In reality, these responses are affected by other variables such as motor noise, task strategies, and attentional lapses. However, these sources of trial-to-trial response variability are difficult to estimate, so here we ignore them for simplicity. For each participant, we found the pair of free parameters that maximized the likelihood of all their heading responses across all trials 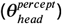.

The likelihood for each trial was computed by reconstructing the optic flow signals presented to the participant, given their gaze direction (*θ*_*eye*_, *α*_*eye*_) and gaze velocity (*ω*_*eye*_), which were measured during each trial using the eye tracker. To ensure that the ideal observer had access to the same motion signals as the human participant, motion signals were only provided at retinal locations corresponding to the experimental display. For trials in which the participant produced multiple smooth pursuits, model responses were generated for every pursuit, and then averaged. Model responses were only generated using gaze data from smooth pursuit phases (ignoring saccadic eye movements, during which visual processing is mostly suppressed).

#### Eccentricity-based downsampling

We implemented and compared three different types of eccentricity-based downsampling in an attempt to incorporate these functional mechanisms into our model. The first model has no downsampling mechanism (as a baseline). The second model downsamples the input by imposing eccentricity-based distance thresholds on the points that comprise the stimulus, using standard visual acuity measurements (Weymouth, 1958). The third and fourth models apply eccentricity-based spatial pooling. Using motion-sensitive receptive field sizes measured by McKee and Nakayama (1984), we tile the visual field with circular receptive fields (ensuring coverage via a minimum overlap of one receptive field radius) that output the linear sum of all points that fall within them. Be-cause it is unknown whether the sensory noise (from Crowell and Banks, 1996) perturbs the motion signal before or after spatial pooling (Crowell and Banks (1996) measured thresholds for single dots), the third model implements the noise before pooling, and the fourth model implements it after pooling. A comparison of the four models reveals that the fourth (noise post pooling) explains the greatest variance in human behavior, although all models performed similarly well (Table 4). This indicates that motion downsampling is somewhat important for maximizing model performance, although not critical. For all reported analyses in the main text, we present data from the best (fourth) model.

**Table 4.** Variance explained by Bayesian ideal observers with different eccentricity-dependent downsampling methods. *R*^2^ and noise-corrected *R*^2^ for different variants of the model, trained and tested via 10-fold cross-validation on data from Experiment 1.

| | $R^2$ | Noise-Corrected $R^2$ |
| --- | --- | --- |
| Baseline (no downsampling) | .15 | .64 |
| Acuity-based separation | .15 | .66 |
| Spatial pooling (noise before pooling) | .15 | .64 |
| Spatial pooling (noise after pooling) | .16 | .71 |

#### Estimating variance explained

Comparing model performance against a baseline is critical for distinguishing between a model that explains human data poorly, and a noisy dataset that cannot be explained without overfitting.

Techniques for estimating baselines for explainable variance typically include presenting the same stimulus multiple times, and measuring participant response consistency. In our experiments, no two stimuli are exactly the same, since they are defined by trial-to-trial gaze dynamics, which cannot be precisely controlled. Here we formulate a method of estimating ceiling performance of our model based on the available data.

Four variables—gaze azimuth (*θ*_*eye*_), gaze elevation (*α*_*eye*_), gain (*g*), and heading direction (*θ*_*head*_))— uniquely define the flow field produced by each trial’s stimulus on the participants’ retina. This is effectively a 4-dimensional vector space where, insofar as these variables predict heading perception, nearby locations produce similar heading responses. Using weighted k-nearest neighbor interpolation, it is possible to predict the response generated by any arbitrary combination of these 4 variables. In this model, *k* is the number of nearest samples (defined by smallest euclidean distance relative to test trial) to average over, and the weights (***w***) scale 3 of the 4 dimensions based on their relative predictive power. Optimal values of *k* and ***w*** for each participant were estimated by minimizing the squared prediction error via leave-one-out cross validation on individual trials (minima identified using Nelder-Mead optimization). Averaged over participants, the optimal 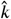 was 13.63 samples, and optimal 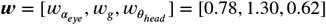. Then, using the optimal parameters, we computed the *R*^2^ between the interpolated responses, and the human data, to obtain participant-specific estimates of the maximum explainable variance. The ratio between the model variance explained, and maximum explainable variance, gives a noise-corrected estimate of the variance explained.

## Acknowledgements

This work was partially funded by Alcon Research LLC.

